# Altering nuclear import in early *Xenopus laevis* embryos affects later development

**DOI:** 10.1101/510313

**Authors:** Predrag Jevtić, Daniel L. Levy

## Abstract

More than just a container for DNA, the nucleus carries out a wide variety of critical and highly regulated cellular functions. One of these functions is nuclear import, and in this study we investigate how altering nuclear import impacts developmental progression and organismal size. During early *Xenopus laevis* embryogenesis, the timing of a key developmental event, the midblastula transition (MBT), is sensitive to nuclear import factor levels. How might altering nuclear import and MBT timing in the early embryo affect downstream development of the organism? We microinjected *X.laevis* two-cell embryos to increase levels of importin α or NTF2, resulting in differential amounts of nuclear import factors in the two halves of the embryo. Compared to controls, these embryos exhibited delayed gastrulation, curved neural plates, and bent tadpoles with different sized eyes. Furthermore, embryos microinjected with NTF2 developed into smaller froglets compared to control microinjected embryos. We propose that altering nuclear import and size affects MBT timing, cell size, and cell number, subsequently disrupting later development. Thus, altering nuclear import early in development can affect function and size at the organismal level.

## INTRODUCTION

More than just a container for DNA, the nucleus carries out a wide variety of critical and highly regulated cellular functions. The nuclear envelope (NE) is composed of a double lipid bilayer. The outer nuclear membrane is continuous with the endoplasmic reticulum while the inner nuclear membrane is lined and supported by the nuclear lamina, composed of a meshwork of lamin intermediate filaments and lamin-associated proteins (1, 2). Nuclear pore complexes (NPC) that mediate nucleocytoplasmic transport are inserted into the NE at sites where the inner and outer nuclear membranes fuse (2–5). After mitosis and nuclear reassembly, lamins are imported into the nucleus along with other proteins containing nuclear localization signals (NLS). Classical nuclear import is mediated by importin α/β karyopherins, which bind NLS-containing proteins and ferry them across the NPC and into the nucleus. Within the nucleus, Ran in its GTP-bound state binds to importin β thereby releasing NLS cargos. Another key player in this process is NTF2, a dedicated nuclear import factor for Ran (6–10). Associated with the NPC, NTF2 has been shown to reduce import of large cargos (11–13). While nuclear import is critical for a wide variety of cell functions (14, 15), in this study we investigate how altering nuclear import impacts developmental progression and organismal size.

In *Xenopus*, levels of two nuclear import factors were shown to tune rates of nuclear import, which coincidentally also impacted nuclear growth. Increased importin α levels generally positively scale with nuclear import and size while increased NTF2 negatively regulates import of large cargos and nuclear size, although differential effects are observed when these factors are present at very high levels and depending on the cellular context (12, 13). Nuclear lamins represent one imported cargo that contributes to nuclear growth (16). How might nuclear import impact development? During early *X.laevis* development, the first twelve cleavage cell divisions occur rapidly with little new transcription occurring (i.e. stages 1-8). Stage 8 coincides with the midblastula transition (MBT) when zygotic transcription is upregulated, cell cycles slow, and there is onset of cell division asynchrony and motility (17–20). While the DNA-to-cytoplasm ratio is one important factor that determines when the MBT initiates (17, 18, 21–23), we previously demonstrated that altering nuclear import and size in early *X.laevis* embryos also affects MBT timing (24, 25). An important question raised by these studies is how altering nuclear import and MBT timing in the early embryo affects downstream development of the organism.

Here we test how altering levels of nuclear import factors in the early embryo affects later development. Previous work has shown how levels of importin α and NTF2 impact nuclear import in *Xenopus* and cell culture, consistent with computer models (12, 13, 26–29). It has also been shown that the levels of NTF2 inversely correlate with nuclear enlargement during melanoma progression and that NTF2 overexpression was sufficient to reduce nuclear size in primary melanoma (13). There is growing evidence that early embryogenesis and cancer progression share similar cellular features, such as rapid cell proliferation and increased cell motility, and that many embryo-specific genes and signaling pathways are reactivated in cancer (30, 31). For these reasons, we were particularly interested to test the developmental consequences of altering NTF2 levels because of its potential involvement in carcinogenesis (13, 32). In this study, we investigate how altering nuclear import in *X.laevis* embryos impacts gastrulation, neurulation, and development of tadpoles and froglets.

## RESULTS AND DISCUSSION

We microinjected one blastomere of two-cell stage embryos with mRNA encoding nuclear import factors along with fluorescent dextran to trace cells that received the mRNA (Fig. 1A). One important advantage of this approach is that the uninjected half of the embryo serves as an internal control, thus facilitating the observation of any developmental differences between the two halves of the embryo. In some cases we differentially microinjected the two blastomeres to maximize potential nuclear import differences in the two halves (i.e. importin α/lamin B3 in one half and NTF2 in the other half). For control experiments, embryos were microinjected with mRNA encoding either GFP or histone H2B-GFP. Embryos were microinjected with mRNA amounts previously shown to maximally alter nuclear size in vivo (Fig. S1) (12, 13, 24).

**Figure 1:**
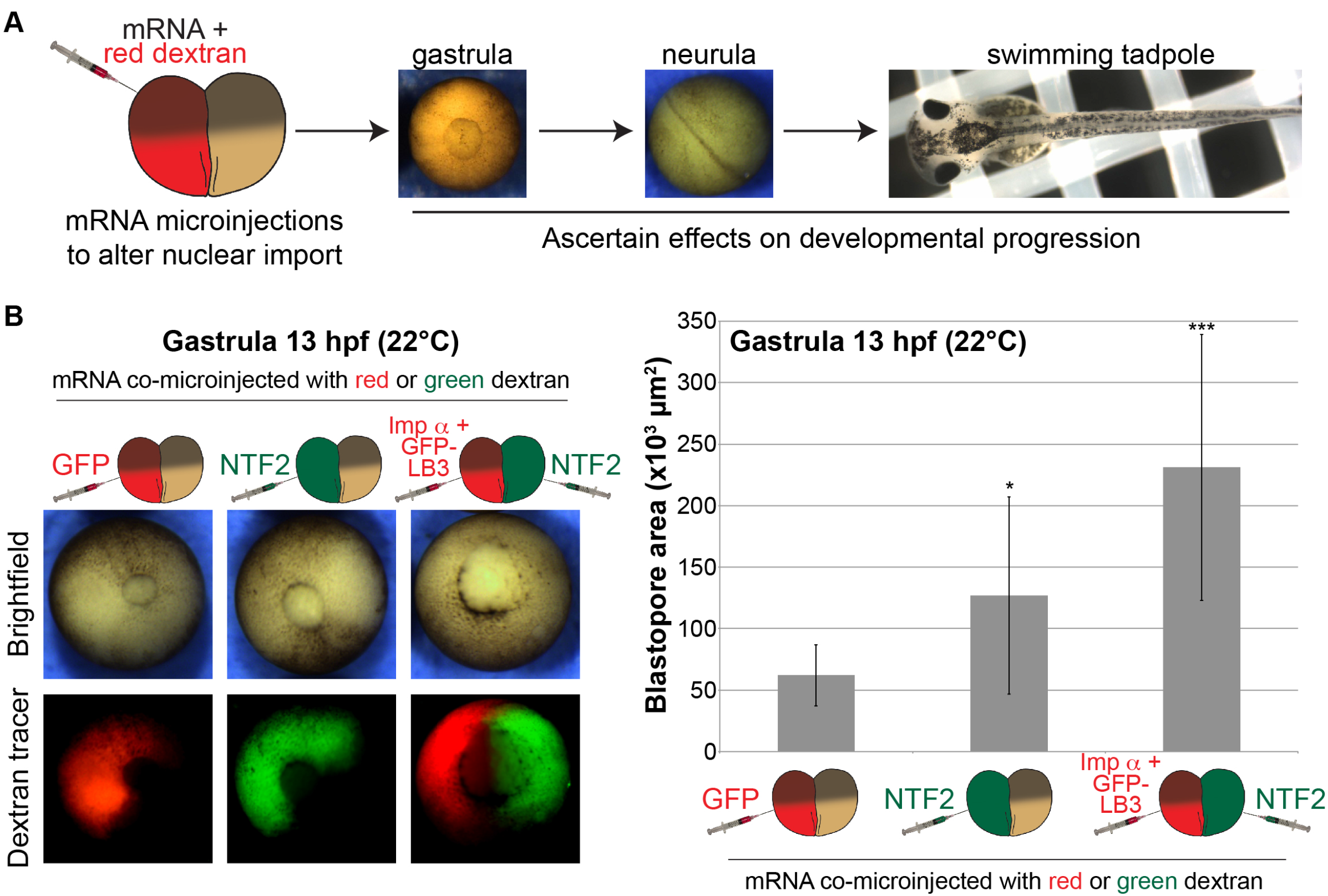
Differential nuclear import in the two halves of an early embryo delays gastrulation. **(A)** Experimental approach. One blastomere of a two-cell stage *X.laevis* embryo was co-microinjected with mRNA to alter nuclear import factor levels and fluorescently labeled dextran as a cell tracer. Embryos were allowed to develop to different stages to assess effects on developmental progression. **(B)** Microinjections were performed as shown in (A) with 250 pg GFP mRNA, 175 pg NTF mRNA, or 250 pg importin α mRNA + 250 pg GFP-LB3 mRNA. These amounts, that maximally affect nuclear size (13, 24), were used in all experiments. Microinjected two-cell embryos were allowed to develop to 13 hpf gastrula. Representative vegetal pole images are shown. Blastopore area was measured and averaged for 10-13 embryos per condition. Error bars represent SD. *** p<0.005, * p<0.05.

To quantify the timing of gastrulation, we measured blastopore size in microinjected embryos 13 hours post fertilization (hpf). We previously showed that increasing importin α and lamin B3 (LB3) expression levels led to premature onset of the MBT and accelerated blastopore closure during gastrulation (24). Conversely, NTF2 microinjection delayed blastopore closure, and an even greater delay was observed when half the embryo was microinjected with NTF2 and the other half was microinjected with importin α/LB3 (Fig. 1B). These data suggest that inducing differential nuclear import in the two halves of the embryo affects the timing of gastrulation.

We next examined how neurulation was affected when nuclear import was manipulated early in development. Altering levels of nuclear import factors in half of the embryo frequently resulted in differential timing of neural plate closure in the two halves of the embryo (Fig. S2A, Videos 1-4). Consequently, these embryos exhibited a curved neural plate (Fig. 2A, S3A, Videos 5-6). This phenotype was observed in over 65% of NTF2-microinjected embryos, increasing to 85% for embryos microinjected to maximize the import differential in the two halves of the embryo (Fig. 2B). Similar phenotypes were observed when nuclear import factor mRNA was co-microinjected with H2B-GFP mRNA instead of dextran tracer and with frogs and embryos derived from two different frog colonies (Fig. S3). Furthermore, microinjection of importin α alone induced a similar effect as importin α/LB3 (Fig. 2 and S3), suggesting the curved neural plate phenotype resulted from altered nuclear import. The formation of curved neural plates was dependent on sufficiently altering nuclear import as microinjecting lower amounts of NTF2 mRNA (e.g. 50 pg) failed to induce neural plate bending (data not shown). When one-cell embryos were microinjected we observed no effect on the morphology of the neural plate compared to controls, demonstrating that the curved phenotype was dependent on there being differential amounts of nuclear import factors in the two halves of the embryo (Fig. S3B).

**Figure 2:**
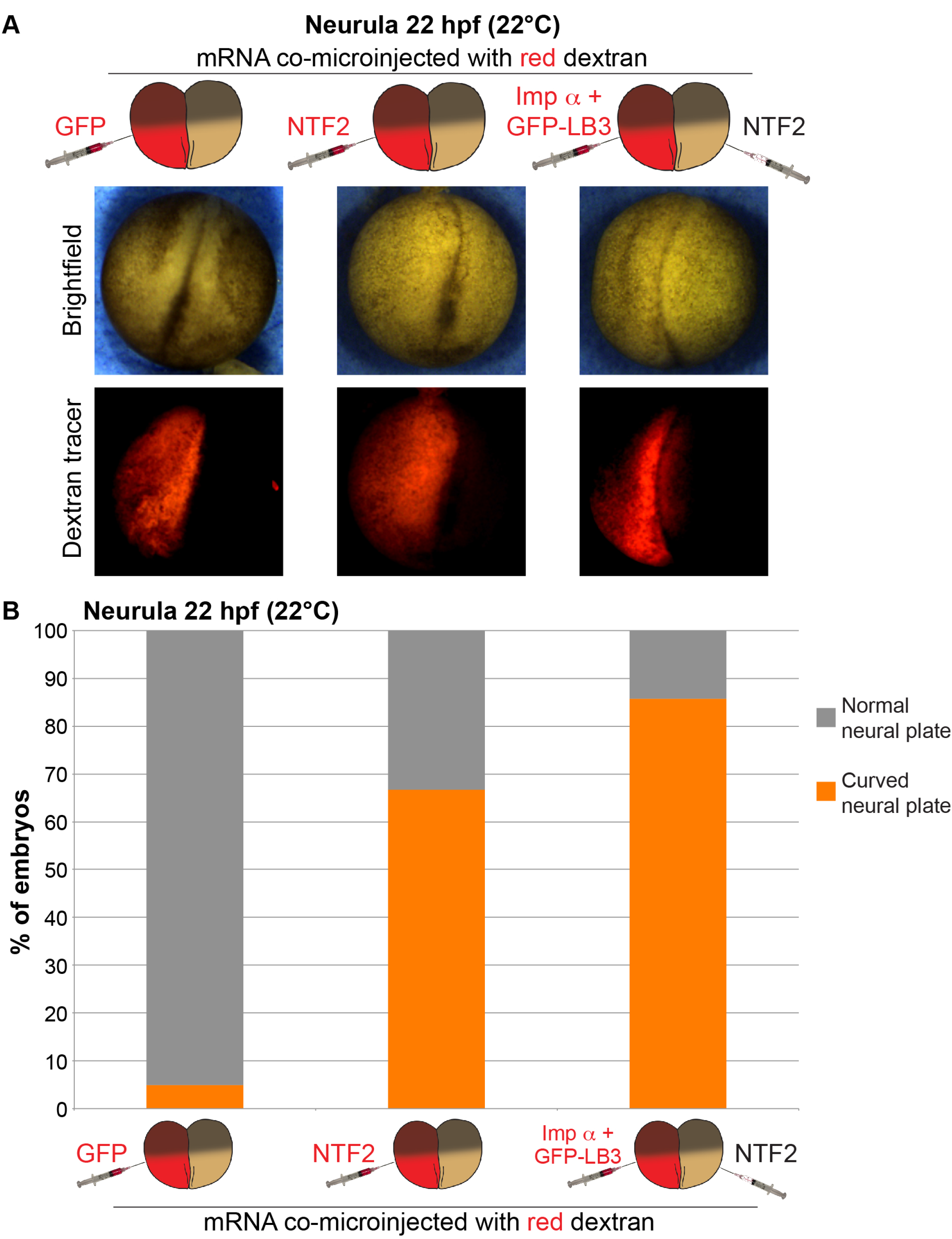
Differential nuclear import in the two halves of an early embryo leads to neural plate curvature. **(A)** Two-cell embryos were microinjected as indicated and allowed to develop to 22 hpf neurula. Representative images are shown. **(B)** Neurula were scored as having normal or curved neural plates by drawing a line through the middle of the embryo. Embryo numbers: n=19 for GFP, n=21 for NTF2, n=7 for imp α + GFP-LB3/NTF2.

Interestingly, the neural plate generally curved toward the NTF2 injected side or away from the importin α/LB3 injected side (Fig. 2A, S3). Altering MBT timing and the onset of longer cell cycles has been shown to indirectly impact cell size (24, 33–35). In particular, in the half of the embryo with increased importin α levels and nuclear size, early onset of longer cell cycles results in larger cells, potentially explaining why the neural plate curved away from that side of the embryo. Indeed, consistent with these previous reports, surface imaging showed smaller cells on the NTF2-injected side and larger cells on the importin α-injected size (Fig. S2B-C).

We next asked if the bent neural plate phenotype was propagated later in development. More than 30% of NTF2-microinjected embryos exhibited a bent tadpole phenotype, again with the bend occurring toward the NTF2-microinjected side (Fig. 3, S4-S5). We also observed more than 30% of tadpoles with a smaller eye on the NTF2-microinjected side, with 16% of embryos showing both the small eye and bent body phenotype (Fig. 3, S4). The small eye phenotype was exacerbated in embryos microinjected to maximize nuclear import differences in the two halves of the embryo (Fig. 3B). Embryos microinjected at the one-cell stage did not develop into bent tadpoles (Fig. S5A), similar to what was observed for neurula. Similar bent tadpoles were observed with frogs and embryos derived from two different frog colonies as well as for embryos microinjected with importin α/LB3 or importin α alone (Fig. 3 and S5).

**Figure 3:**
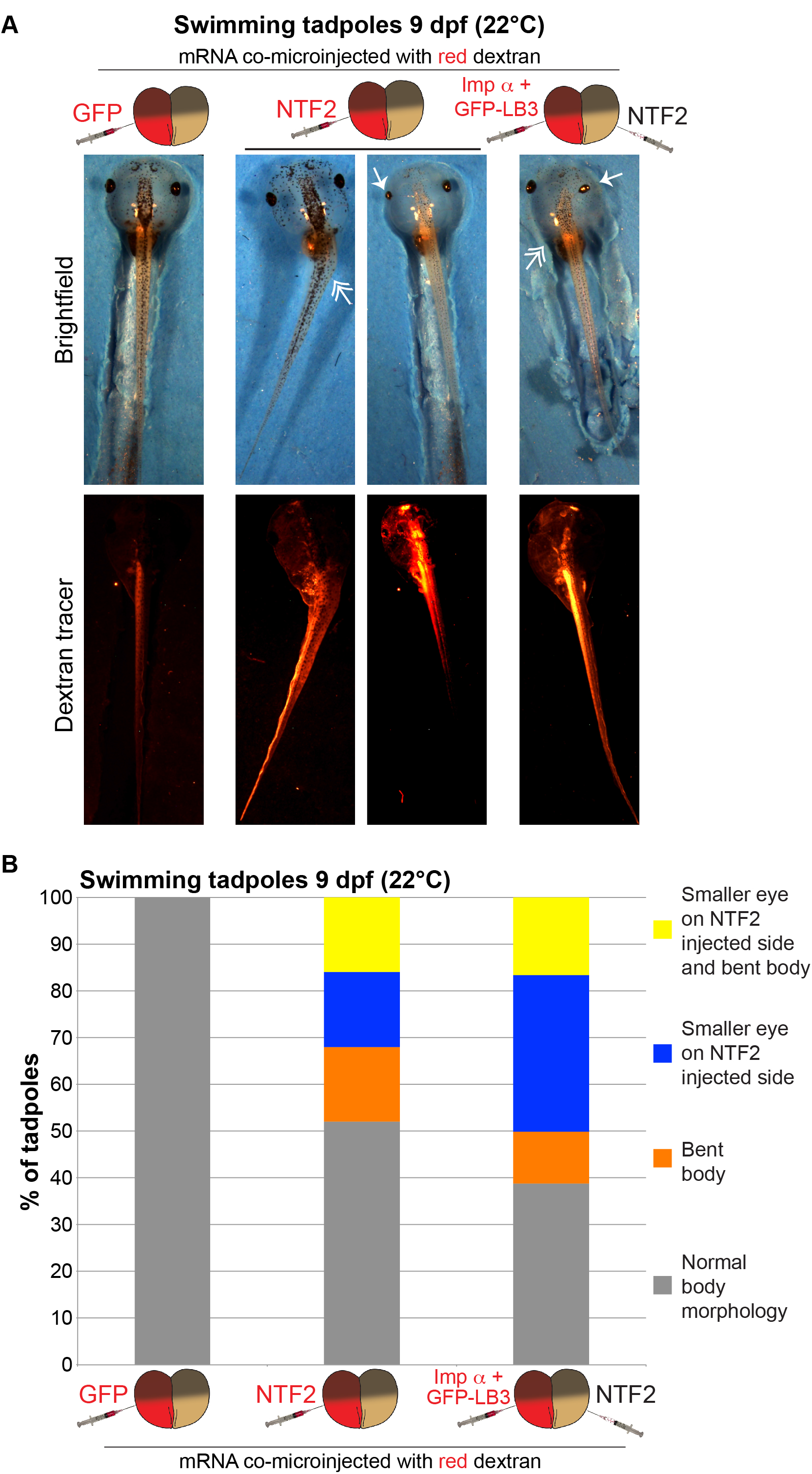
Differential nuclear import in the two halves of an early embryo leads to bent tadpoles. **(A)** Two-cell embryos were microinjected as indicated and allowed to develop into 9 dpf swimming tadpoles. Representative images are shown. Singleheaded arrows indicate small eyes. Double-headed arrows indicate bent bodies. **(B)** Tadpoles were scored as indicated by measuring eye areas and body axis angles. Embryo numbers: n=10 for GFP, n=25 for NTF2, n=18 for imp α + GFP-LB3/NTF2.

Lastly, we allowed microinjected embryos to develop into 4-month-old froglets. NTF2-microinjected embryos gave rise to significantly smaller froglets, with a small proportion exhibiting defective body morphologies (Fig. 4). Many of the NTF2-injected froglets did not survive to adulthood, however those that did were smaller than their control sexed counterparts (Fig. S6A). It is possible that the frogs generated from NTF2-injected embryos were smaller due to malnutrition associated with their morphological defects. Eggs produced by these smaller females were the same size as controls (Fig. S6B), although nuclei assembled de novo in extract isolated from their eggs were smaller (data not shown). Interestingly, erythrocyte nuclei were smaller in animals derived from NTF2-injected embryos compared to sexed controls (Fig. S6C). Consistent with this coordination between body size and erythrocyte nuclear size, male erythrocyte nuclei were smaller than those in females, as has been observed in other amphibian species where males are smaller than females (36, 37). The erythrocyte nuclear-to-cytoplasmic volume ratio was the same for control females and males but reduced in the case of NTF2 microinjection (Table S1).

**Figure 4:**
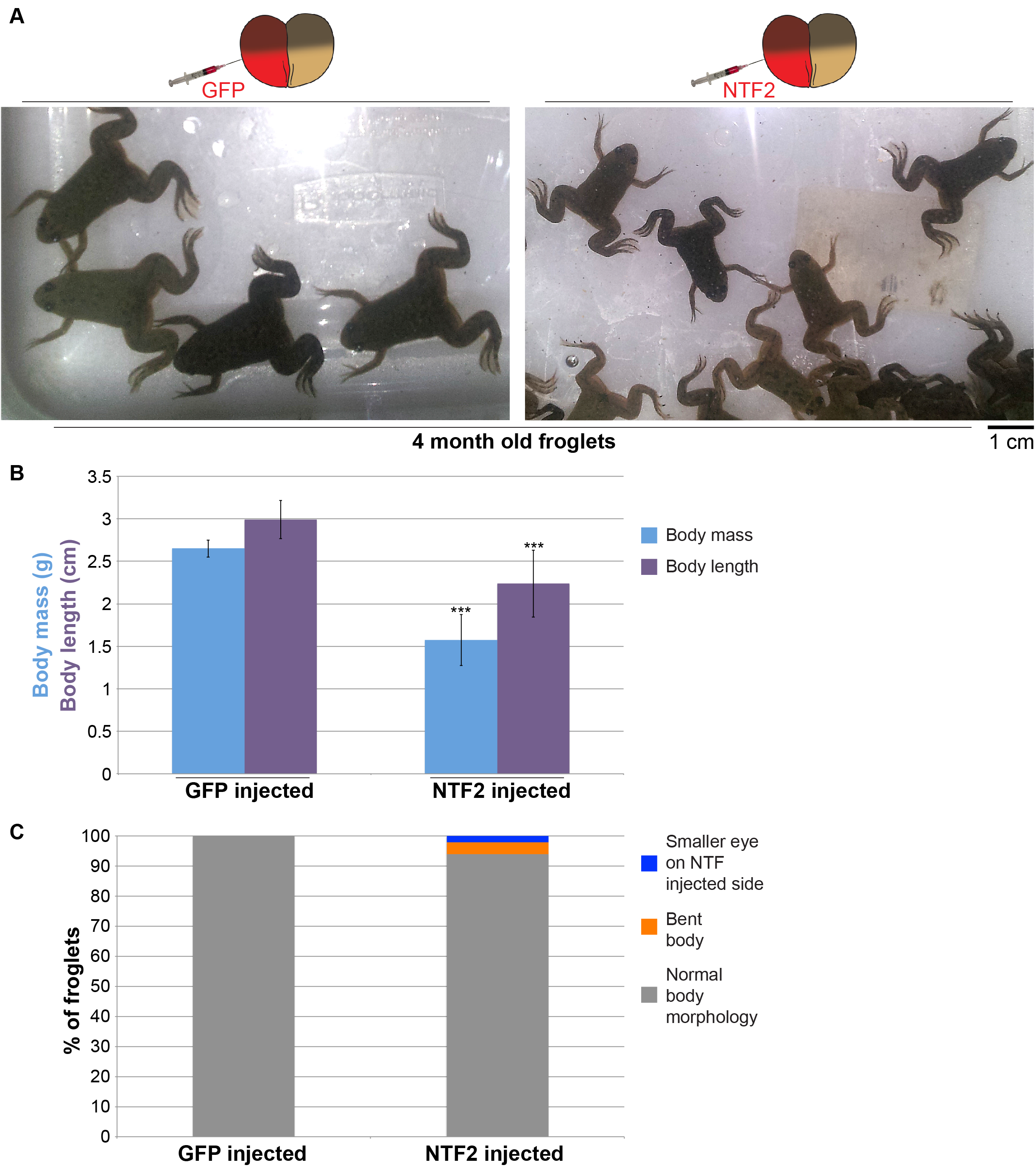
Differential nuclear import in the two halves of an early embryo leads to smaller froglets. Two-cell embryos were microinjected as indicated and allowed to develop into 4-month-old froglets. Froglet numbers: n=33 for GFP and n=50 for NTF2. **(A)** Representative froglets. **(B)** Quantification of froglet body mass and length. **(C)** Scoring of froglets with altered body morphology as indicated. Error bars represent SD. *** p<0.005.

Taken together, we show that altering the levels of nuclear import factors in the early embryo leads to downstream effects on gastrulation, neurulation, and the development of tadpoles and froglets. Specifically, NTF2, importin α, and lamin expression levels impact developmental outcomes. These results show how altering nuclear import can, perhaps indirectly, affect function and size at the organismal level. Furthermore, given the defects we observed in neural plate and body morphologies, our findings may be relevant to a wide range of diseases associated with neural tube defects (38). Because nuclear import impinges on a variety of different cellular functions (14, 15), we cannot at this time specify which altered function might be responsible for the observed effects on development. We will note that importin α/LB3 microinjection increases nuclear size while NTF2 microinjection decreases nuclear size (Fig. S1) (12,13, 24). One possibility is that altering nuclear import and size in the early embryo leads to changes in MBT timing that in turn impact cell size and number, subsequently disrupting later stages of development.

## MATERIALS AND METHODS

### Plasmids

Plasmids consisting of pCS2+ containing the coding sequences for human importin α2-E (pDL17), *X.tropicalis* GFP-LB3 (pDL19), and NTF2 (pDL18) were described previously (12, 13). For control injections, we used GFP mRNA expressed from pCS107-GFP3STOP or H2B-GFP mRNA expressed from CS2-H2BeGFP (gifts from John Wallingford, University of Texas at Austin).

### *Xenopus laevis* embryos and microinjections

*X. laevis* embryos were obtained by in vitro fertilization of freshly laid *X. laevis* eggs with crushed *X. laevis* testes (39). Only batches with greater than 90% fertilization efficiency were used. Twenty minutes after fertilization, embryos were de-jellied in 2.5% cysteine pH 7.8 dissolved in 1/3x MMR (20x MMR = 2 mM EDTA, 2 M NaCl, 40 mM KCl, 20 mM MgCl_2_, 40 mM CaCl_2_, 100 mM HEPES pH 7.8). Embryos were staged according to (40). All *Xenopus* procedures and studies were conducted in compliance with the US Department of Health and Human Services Guide for the Care and Use of Laboratory Animals. Protocols were approved by the University of Wyoming Institutional Animal Care and Use Committee (Assurance # A-3216-01).

Following linearization of pCS107-GFP3STOP, CS2-H2BeGFP, pDL17, pDL18, and pDL19, mRNA was expressed from the SP6 promoter using the mMessage mMachine kit (Ambion). Embryos at the one-cell or two-cell stage were transferred to 1/3 MMR plus 2.5% Ficoll and microinjected with 10 nL volumes using a PicoSpritzer III (Parker). Different amounts of mRNA were injected by varying the concentration of the mRNA stock solution. Unless otherwise indicated, the following mRNA amounts were used for each microinjection: 250 pg GFP, 100 pg H2B-GFP, 175 pg NTF2, 250 pg importin α, 250 pg GFP-LB3. This amount of NTF2 mRNA maximally decreases nuclear size (13), while these amounts of importin α and GFP-LB3 mRNA maximally increase nuclear size (24). After 45 minutes, the buffer was changed to 1/3x MMR and embryos were allowed to develop to desired stages. Tadpoles were grown in 1/3x MMR in small tanks with water filtration at room temperature. During metamorphosis, froglets were grown in the same water as adults at room temperature. Tadpoles and froglets were fed tadpole frog brittle (Nasco SA05964) and post-metamorphic frog brittle (Nasco SB29027), respectively.

In most experiments, one blastomere of a two-cell embryo was co-microinjected with mRNA and 50 ng of a fluorescently labeled dextran that served as a marker for the injected half. Dextrans used were lysine-fixable tetramethylrhodamine-labeled dextran, 70,000 MW (ThermoFisher, D1818) or lysine-fixable fluorescein-labeled dextran, 70,000 MW (ThermoFisher, D1822). For control experiments, mRNA expressing GFP or H2B-GFP was used.

### Microscopy and image quantification

For Figure S1, microinjected embryos at stage 11 were transferred to 1/3x MMR containing 10 μg/ml Hoechst. Subsequently, embryos were squashed between a glass coverslip and slide for imaging. For erythrocyte measurements, adult frogs were anesthetized in 0.05% benzocaine prior to blood draw. Abdominal skin was dried and punctured with a sterile 30G½ needle. Blood was immediately smeared on the surface of a glass slide and fixed in methanol for 3 minutes at room temperature. Blood smears were stained in 1x Giemsa stain (Sigma G5637) for 45 minutes at room temperature. Nuclei and erythrocytes were visualized with an Olympus BX51 fluorescence microscope using an Olympus UPLFLN 20x (N.A. 0.50, air) objective. Images were acquired with a QIClick Digital CCD Camera, Mono, 12-bit (model QIClick-F-M-12) at room temperature using Olympus cellSens software. Nuclear and erythrocyte crosssectional areas were quantified from original thresholded images using cellSens Dimension imaging software (Olympus).

Brightfield and fluorescence imaging of eggs, embryos, and tadpoles was performed with an Olympus SZX16 research fluorescence stereomicroscope, equipped with Olympus DP72 camera, 11.5x zoom microscope body, and SDFPLAPO1XPF objective. Brightfield time-lapse imaging of embryos was performed at room temperature, and images were acquired every 5 minutes. Discontinuous light was used to illuminate embryos, controlled with a digital adjustable cycle timer (CT-1 Short Cycle Timer, Innovative Grower Corp). Swimming tadpoles were anesthetized in 1/3x MMR containing 0.05% benzocaine prior to imaging. Blastopore area and egg size were quantified from original thresholded images using cellSens Dimension imaging software (Olympus). Eye areas and body angles in swimming tadpoles were measured from original images using cellSens Dimension imaging software measurement tools (Olympus). Froglets were anesthetized in 0.05% benzocaine to measure body mass and length. Froglets and frogs were imaged in plastic containers using a cell phone camera.

Where indicated, confocal imaging was performed on a spinning-disk confocal microscope based on an Olympus IX71 microscope stand equipped with a five line LMM5 laser launch (Spectral Applied Research) and Yokogawa CSU-X1 spinning-disk head. Confocal images were acquired with an EM-CCD camera (ImagEM, Hamamatsu). Z-axis focus was controlled using a piezo Pi-Foc (Physik Instrumentes), and multiposition imaging was achieved using a motorized Ludl stage. An Olympus UPLSAPO 20xOЮ.85na objective was used. Image acquisition and all system components were controlled using Metamorph software.

### Statistics

Averaging and statistical analysis were performed for independently repeated experiments. Two-tailed Student’s t tests assuming equal variances were performed in Excel (Microsoft) to evaluate statistical significance. The p values, sample sizes, and error bars are given in the figure legends.

## ACKNOWLEDGMENTS

We thank John Wallingford (University of Texas, Austin) for plasmids, Rebecca Heald (University of California, Berkeley) for support in the early stages of this research, and Karen White (University of Wyoming) for critical reading of the manuscript.

## Abbreviations

NE: nuclear envelope
MBT: midblastula transition
hpf: hours post fertilization
dpf: days post fertilization
NTF2: nuclear transport factor 2
LB3: lamin B3
imp α: importin α
NLS: nuclear localization signal
NPC: nuclear pore complex

## REFERENCES

1. Dittmer TA, Misteli T. The lamin protein family. Genome Biol. 2011;12(5):222.

2. Wilson KL, Berk JM. The nuclear envelope at a glance. J Cell Sci. 2010;123(Pt 12):1973–8.

3. Rothballer A, Kutay U. SnapShot: the nuclear envelope II. Cell. 2012;150(5):1084–e1.

4. Rothballer A, Kutay U. SnapShot: The nuclear envelope I. Cell. 2012;150(4):868–e1.

5. Misteli T, Spector DL. The Nucleus. Cold Spring Harbor, New York: Cold Spring Harbor Laboratory Press; 2011. 517 p.

6. Madrid AS, Weis K. Nuclear transport is becoming crystal clear. Chromosoma. 2006; 115(2):98–109.

7. Stewart M. Molecular mechanism of the nuclear protein import cycle. Nat Rev Mol Cell Biol. 2007;8(3):195–208.

8. Fried H, Kutay U. Nucleocytoplasmic transport: taking an inventory. Cell Mol Life Sci. 2003;60(8): 1659–88.

9. Dickmanns A, Kehlenbach RH, Fahrenkrog B. Nuclear Pore Complexes and Nucleocytoplasmic Transport: From Structure to Function to Disease. Int Rev Cell Mol Biol. 2015;320:171–233.

10. Hutten S, Kehlenbach RH. CRM1-mediated nuclear export: to the pore and beyond. Trends Cell Biol. 2007;17(4):193–201.

11. Feldherr C, Akin D, Moore MS. The nuclear import factor p10 regulates the functional size of the nuclear pore complex during oogenesis. J Cell Sci. 1998; 111 (Pt 13):1889–96.

12. Levy DL, Heald R. Nuclear size is regulated by importin alpha and Ntf2 in Xenopus. Cell. 2010;143(2):288–98.

13. Vukovic LD, Jevtic P, Zhang Z, Stohr BA, Levy DL. Nuclear size is sensitive to NTF2 protein levels in a manner dependent on Ran binding. J Cell Sci. 2016;129(6):1115–27.

14. Macara IG. Transport into and out of the nucleus. Microbiol Mol Biol Rev. 2001; 65(4):570–94, table of contents.

15. Mackmull MT, Klaus B, Heinze I, Chokkalingam M, Beyer A, Russell RB, et al. Landscape of nuclear transport receptor cargo specificity. Mol Syst Biol. 2017;13(12):962.

16. Jevtic P, Edens LJ, Li X, Nguyen T, Chen P, Levy DL. Concentration-dependent Effects of Nuclear Lamins on Nuclear Size in Xenopus and Mammalian Cells. J Biol Chem. 2015;290(46):27557–71.

17. Newport J, Kirschner M. A major developmental transition in early Xenopus embryos: I. characterization and timing of cellular changes at the midblastula stage. Cell. 1982;30(3):675–86.

18. Newport J, Kirschner M. A major developmental transition in early Xenopus embryos: II. Control of the onset of transcription. Cell. 1982;30(3):687–96.

19. Newport JW, Kirschner MW. Regulation of the cell cycle during early Xenopus development. Cell. 1984;37(3):731–42.

20. Collart C, Owens ND, Bhaw-Rosun L, Cooper B, De Domenico E, Patrushev I, et al. High-resolution analysis of gene activity during the Xenopus mid-blastula transition. Development. 2014;141(9):1927–39.

21. Clute P, Masui Y. Regulation of the appearance of division asynchrony and microtubule-dependent chromosome cycles in Xenopus laevis embryos. Dev Biol. 1995; 171 (2):273–85.

22. Kobayakawa Y, Kubota HY. Temporal pattern of cleavage and the onset of gastrulation in amphibian embryos developed from eggs with the reduced cytoplasm. J Embryol Exp Morphol. 1981;62:83–94.

23. Clute P, Masui Y. Microtubule dependence of chromosome cycles in Xenopus laevis blastomeres under the influence of a DNA synthesis inhibitor, aphidicolin. Dev Biol. 1997;185(1):1–13.

24. Jevtic P, Levy DL. Nuclear size scaling during Xenopus early development contributes to midblastula transition timing. Curr Biol. 2015;25(1):45–52.

25. Jevtic P, Levy DL. Both Nuclear Size and DNA Amount Contribute to Midblastula Transition Timing in Xenopus laevis. Sci Rep. 2017;7(1):7908.

26. Gorlich D, Seewald MJ, Ribbeck K. Characterization of Ran-driven cargo transport and the RanGTPase system by kinetic measurements and computer simulation. Embo J. 2003;22(5):1088–100.

27. Riddick G, Macara IG. A systems analysis of importin-{alpha}-{beta} mediated nuclear protein import. J Cell Biol. 2005;168(7):1027–38.

28. Riddick G, Macara IG. The adapter importin-alpha provides flexible control of nuclear import at the expense of efficiency. Mol Syst Biol. 2007;3:118.

29. Smith AE, Slepchenko BM, Schaff JC, Loew LM, Macara IG. Systems analysis of Ran transport. Science. 2002;295(5554):488–91.

30. Ma Y, Zhang P, Wang F, Yang J, Yang Z, Qin H. The relationship between early embryo development and tumourigenesis. J Cell Mol Med. 2010;14(12):2697–701.

31. Aiello NM, Stanger BZ. Echoes of the embryo: using the developmental biology toolkit to study cancer. Dis Model Mech. 2016;9(2):105–14.

32. Jevtic P, Levy DL. Mechanisms of nuclear size regulation in model systems and cancer. Adv Exp Med Biol. 2014;773:537–69.

33. Amodeo AA, Jukam D, Straight AF, Skotheim JM. Histone titration against the genome sets the DNA-to-cytoplasm threshold for the Xenopus midblastula transition. Proc Natl Acad Sci U S A. 2015;112(10):E1086–95.

34. Collart C, Allen GE, Bradshaw CR, Smith JC, Zegerman P. Titration of four replication factors is essential for the Xenopus laevis midblastula transition. Science. 2013;341 (6148):893–6.

35. Wang P, Hayden S, Masui Y. Transition of the blastomere cell cycle from cell size-independent to size-dependent control at the midblastula stage in Xenopus laevis. J Exp Zool. 2000;287(2): 128–44.

36. Hota J, Das M, Mahapatra PK. Blood Cell Profile of the Developing Tadpoles and Adults of the Ornate Frog, Microhyla ornata (Anura: Microhylidae). International Journal of Zoology. 2013;2013:716183.

37. Das M, Mahapatra PK. Hematology of Wild Caught Dubois’s Tree Frog Polypedates teraiensis, Dubois, 1986 (Anura: Rhacophoridae). The Scientific World Journal. 2014;2014:491415.

38. Copp AJ, Greene ND. Neural tube defects– –disorders of neurulation and related embryonic processes. Wiley Interdiscip Rev Dev Biol. 2013;2(2):213–27.

39. Sive HL, Grainger RM, Harland RM. Early development of Xenopus laevis: a laboratory manual. Cold Spring Harbor, N.Y.: Cold Spring Harbor Laboratory Press; 2000. ix, 338 p. p.

40. Nieuwkoop PD, Faber J. Normal Table of Xenopus laevis (Daudin). 2nd ed. Amsterdam: North-Holland Publishing Company; 1967.

