## Supplemental Information for "Altering nuclear import in early *Xenopus laevis* embryos affects later development"

Predrag Jevtić, Daniel L. Levy*
Department of Molecular Biology, University of Wyoming, Laramie, WY, 82071

*Corresponding author:


#### SUPPLEMENTARY FIGURE LEGENDS

**Figure S1: Effects of NTF2 and importin α/lamin B3 microinjection on nuclear size.** We previously demonstrated that microinjection of *X. laevis* single-cell embryos with mRNA encoding NTF2 alone or importin α + GFP-lamin B3 resulted in altered nuclear size in stage 8 embryos. After testing a range of mRNA amounts, we determined that 350 pg of NTF2 mRNA maximally decreased nuclear size (1) while co-microinjection of 500 pg each of importin α and GFP-LB3 mRNA maximally increased nuclear size (2, 3). To confirm that these nuclear size effects were detectable in gastrula stage embryos, we microinjected half of these mRNA amounts into one blastomere of two-cell stage embryos, allowed the embryos to develop to stage 11, and quantified nuclear sizes. **(A)** Two-cell embryos were microinjected as indicated with 250 pg GFP mRNA, 175 pg NTF mRNA, or 250 pg importin $\alpha$ mRNA + 250 pg GFP-LB3 mRNA and allowed to develop to 11 hpf gastrula. Embryos were stained with Hoechst. Representative images are shown. **(B)** Nuclei in dextran-injected cells on the embryo surface were imaged and nuclear cross-sectional areas were quantified. For each condition, 10-20 embryos were analyzed. For each condition, 68-72 nuclei were quantified. Error bars represent SD. *** $p<0.005$. Compared to cells that received GFP mRNA (250 pg), NTF2 mRNA microinjection (175 pg) decreased nuclear area by 31% and importin $\alpha$ + GFP-LB3 mRNA co-microinjection (250 pg each) increased nuclear area by 38%. These amounts of mRNA were therefore used throughout the rest of this study.

**Figure S2: Differential nuclear import in the two halves of an early embryo leads to asymmetric neural plate closure and cell size differences. (A)** Two-cell embryos were microinjected as indicated and allowed to develop to 20 hpf neurula. Representative images are shown from Videos 1-4. Note that the dextran images were acquired at the beginning of the time-lapse while the brightfield images were selected later in the time-lapse to highlight asymmetric neural plate closure. For this reason, the brightfield and dextran images do not perfectly align, with the dextran image simply showing the side of the embryo that was microinjected. For the NTF2 microinjection image, a still frame was selected that shows delayed neural plate closure on the microinjected side, however bending of the neural plate toward the microinjected side does not become apparent until later in the time-lapse (see Video 2). **(B)** Two-cell embryos were microinjected as indicated and allowed to develop to stage 8.

Representative confocal embryo surface images are shown. **(C)** Microinjections were performed as indicated and embryos were allowed to develop to 24 hpf. Representative confocal embryo surface images are shown.

**Figure S3: Differential nuclear import in the two halves of an early embryo leads to neural plate curvature. (A)** Two-cell embryos were microinjected as indicated and allowed to develop to 22 hpf neurula. Representative images are shown. **(B)** One blastomere of a two-cell embryo was microinjected with importin $\alpha$ + GFP-LB3 as indicated in the first column. One-cell embryos were microinjected with NTF2 or importin $\alpha$ + GFP-LB3 as indicated in the second and third columns, respectively. Embryos were allowed to develop to 22 hpf neurula. Representative images are shown. Neurula were scored as having normal or curved neural plates by drawing a line through the middle of the embryo. Embryo numbers: n=19 for GFP, n=22 for imp $\alpha$ + GFP-LB3 injected into one cell at 2-cell stage, n=10 for NTF2 injected at the 1-cell stage, n=39 for imp $\alpha$ + GFP-LB3 injected at the 1-cell stage. The GFP microinjection quantification is the same as shown in Fig. 2B. Data presented in Fig. 2/S3B and Fig. S3A were generated from two different frog colonies.

**Figure S4: Quantifying defects in tadpole body morphology. (A)** Two-cell embryos were microinjected as indicated and allowed to develop into 9 dpf swimming tadpoles. Representative images are shown. Eye areas were measured from brightfield images, as shown in red text. Body axis bend angles were measured from brightfield images, as shown in green text. Note that the bent tadpole microinjected with NTF2 is the same one shown in Fig. 3A. **(B)** For each tadpole, the area of the smaller eye was divided by the area of the larger eye to obtain the eye size ratio. Average ratios are plotted for 3-7 tadpoles per condition. **(C)** Average body axis angles are plotted for 4-5 tadpoles per condition. Error bars represent SD. *** $p<0.005$, ** $p<0.01$, * $p<0.05$.

**Figure S5: Differential nuclear import in the two halves of an early embryo leads to bent tadpoles. (A)** Column 1: uninjected embryo. Column 2: One-cell embryo was microinjected with NTF2. Column 3: One blastomere of a two-cell embryo was microinjected with NTF2. Column 4: One-cell embryo was microinjected with importin $\alpha$. Column 5: One blastomere of a two-cell embryo was microinjected with importin $\alpha$. Embryos were allowed to develop into 5-6 dpf swimming tadpoles. Representative images are shown. Double-headed arrows indicate bent bodies. **(B)** Two-cell embryos were microinjected as indicated and allowed to develop into 9 dpf swimming tadpoles. Tadpoles were scored as indicated by measuring body axis angle. Embryo numbers: n=10 for GFP, n=10 for imp $\alpha$ + GFP-LB3. The GFP microinjection quantification is the same as shown in Fig. 3B. Data presented in Fig. 3/S5B and Fig. S5A were generated from two different frog colonies.

**Figure S6: Effects of NTF2 microinjection on frog, egg, and erythrocyte size. (A)** One blastomere of a two-cell embryo was microinjected with NTF2 and embryos were allowed to develop into four-year-old frogs. Single-headed arrow indicates small/absent eye. Double-headed arrow indicates bent body. Frog masses are indicated in parentheses. **(B)** Eggs were collected from the control and NTF2 injected females shown in (A), and egg diameters were measured. Representative images are shown. Egg numbers quantified: n=132 for control, n=46 for NTF2. **(C)** Blood was collected from the animals shown in (A), and erythrocytes were stained with Giemsa. Representative images are shown. Nuclear cross-sectional areas were quantified and averaged. 105-193 nuclei were quantified per condition. Also see Table S1. Error bars represent SD. *** p<0.005, NS not significant.

**Table S1: Erythrocyte measurements**

| | **Female** | **Male** |
|---|---|---|
| **Cell area ($\mu m^2$)** | | |
| **Control** | 162 ± 13 | 142 ± 2 |
| **NTF2-microinjected** | 155 ± 10 | 156 ± 24 |
| **Nuclear area ($\mu m^2$)** | | |
| **Control** | 20.9 ± 3.7 | 18.1 ± 3.9 |
| **NTF2-microinjected** | 18.7 ± 2.9 | 16.3 ± 2.0 |
| **N/C volume ratio (%)** | | |
| **Control** | 4.9 ± 0.6 | 4.8 ± 0.3 |
| **NTF2-microinjected** | 4.3 ± 0.2 | 3.5 ± 0.03 |

Data presented as average ± SD. 120-354 nuclei were quantified per condition. 38-166 cells were quantified per condition.

**VIDEO LEGENDS**

**Video 1: GFP control neural plate closure.** A two-cell embryo was microinjected in the left side with GFP mRNA and allowed to develop to a 20 hpf neurula at 22°C. Brightfield imaging was performed at 5 minute intervals. The total length of the time-lapse is 4 hours.

**Video 2: NTF2 microinjection neural plate closure.** A two-cell embryo was microinjected in the left side with NTF2 mRNA and allowed to develop to a 20 hpf neurula at 22°C. Brightfield imaging was performed at 5 minute intervals. The total length of the time-lapse is 3 hours.

**Video 3: Importin α + GFP-LB3 microinjection neural plate closure.** A two-cell embryo was microinjected in the left side with importin α + GFP-LB3 mRNA and allowed to develop to a 20 hpf neurula at 22°C. Brightfield imaging was performed at 5 minute intervals. The total length of the time-lapse is 5.5 hours.

**Video 4: Importin α + GFP-LB3 versus NTF2 microinjection neural plate closure.** A two-cell embryo was microinjected with importin α + GFP-LB3 mRNA in the left side and with NTF2 mRNA in the right side. The embryo was allowed to develop to a 20 hpf neurula at 22°C. Brightfield imaging was performed at 5 minute intervals. The total length of the time-lapse is 9 hours.

**Video 5: Importin α + GFP-LB3 versus NTF2 microinjection neurula.** A two-cell embryo was microinjected with importin $\alpha$ + GFP-LB3 mRNA in the left side and with NTF2 mRNA in the right side. The embryo was allowed to develop to a 22 hpf neurula at 22°C. Brightfield imaging was performed at 5 minute intervals. The total length of the time-lapse is 3 hours.

**Video 6: NTF2 microinjection neurula.** For the top two embryos, one blastomere of a two-cell embryo was co-microinjected with red dextran and H2B-GFP mRNA. For the bottom two embryos, one blastomere of a two-cell embryo was co-microinjected with NTF2 mRNA, red dextran, and H2B-GFP mRNA. Embryos were allowed to develop to 22 hpf neurula at 22°C. Brightfield imaging was performed at 5 minute intervals. Red and green fluorescence images at the beginning and end of the movie show the sides of the embryos that were microinjected. The total length of the time-lapse is 4.5 hours.

**REFERENCES**

1. Vukovic LD, Jevtic P, Zhang Z, Stohr BA, Levy DL. Nuclear size is sensitive to NTF2 protein levels in a manner dependent on Ran binding. J Cell Sci. 2016;129(6):1115-27.
2. Jevtic P, Levy DL. Nuclear size scaling during Xenopus early development contributes to midblastula transition timing. Curr Biol. 2015;25(1):45-52.
3. Levy DL, Heald R. Nuclear size is regulated by importin alpha and Ntf2 in Xenopus. Cell. 2010;143(2):288-98.

Figure S1

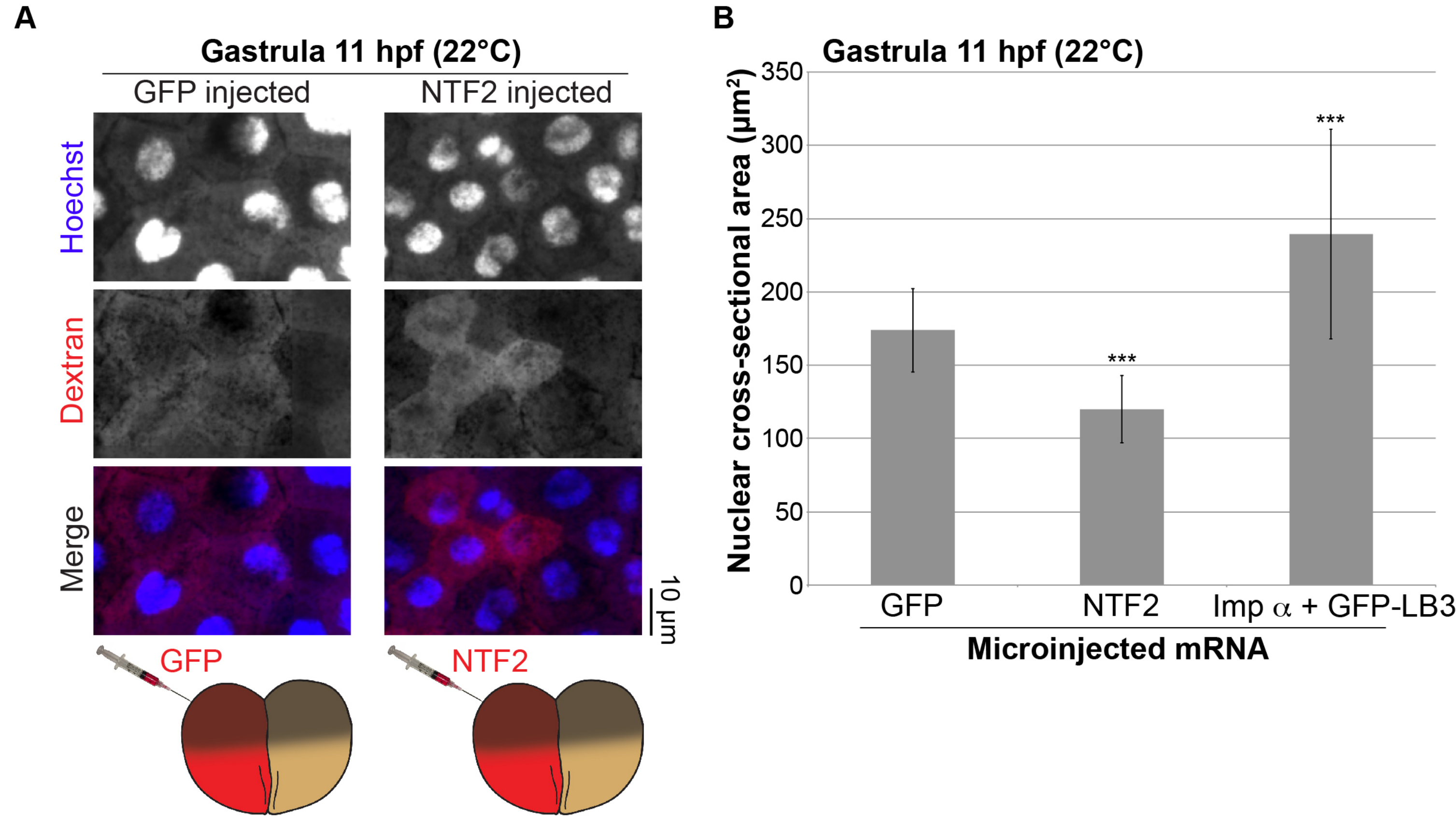
A
Gastrula 11 hpf (22°C)
GFP injected
NTF2 injected
Hoechst
Dextran
Merge
10 µm
GFP
NTF2
B
Gastrula 11 hpf (22°C)
Nuclear cross-sectional area (µm²)
350
300
250
200
150
100
50
0
***
***
GFP
NTF2
Imp α + GFP-LB3
Microinjected mRNA

Figure S2

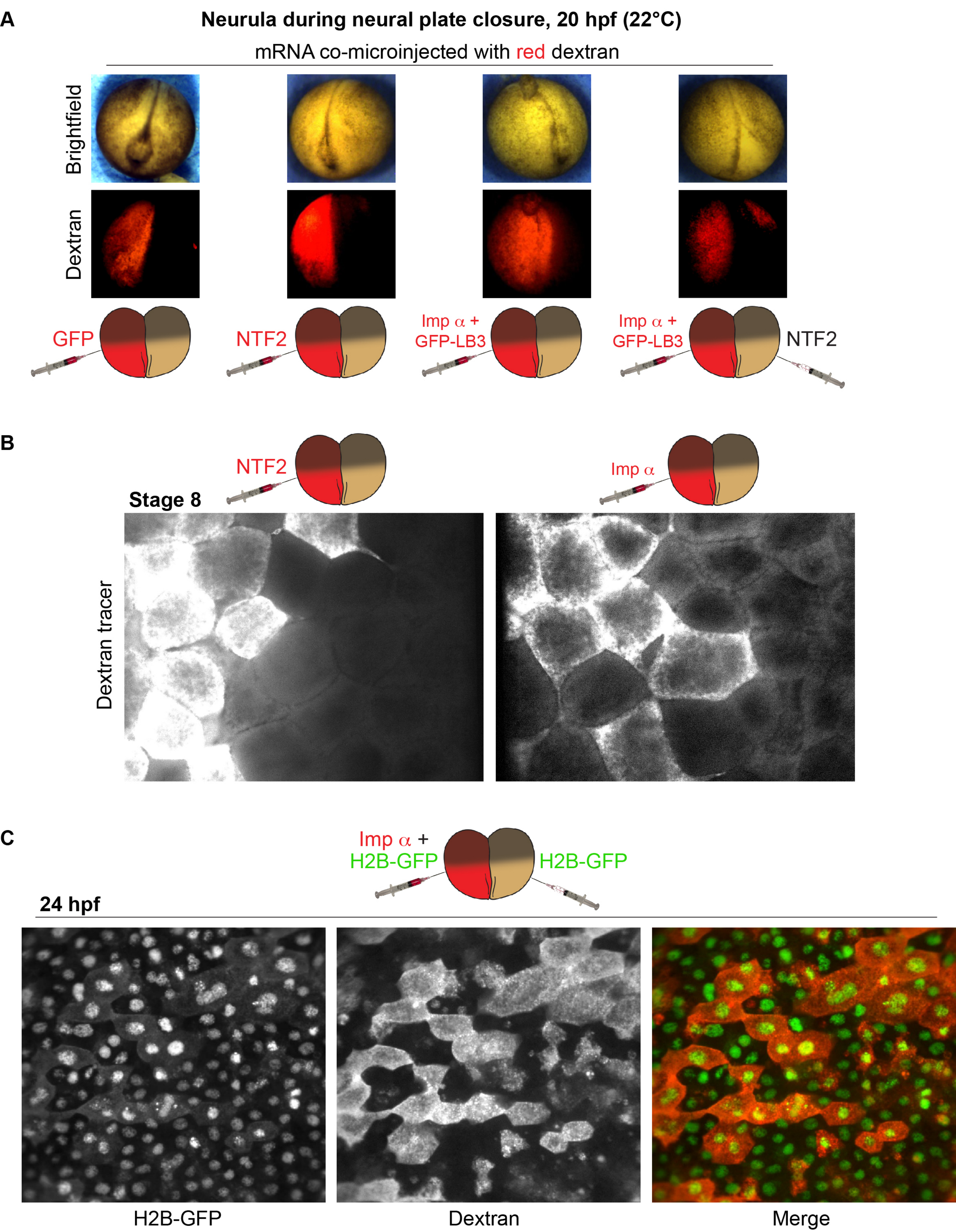
A
Neurula during neural plate closure, 20 hpf (22°C)
mRNA co-microinjected with red dextran
Brightfield
Dextran
GFP
NTF2
Imp α +
GFP-LB3
Imp α +
GFP-LB3
NTF2
B
NTF2
Imp α
Stage 8
Dextran tracer
C
Imp α +
H2B-GFP
H2B-GFP
24 hpf
H2B-GFP
Dextran
Merge

**Figure S3**

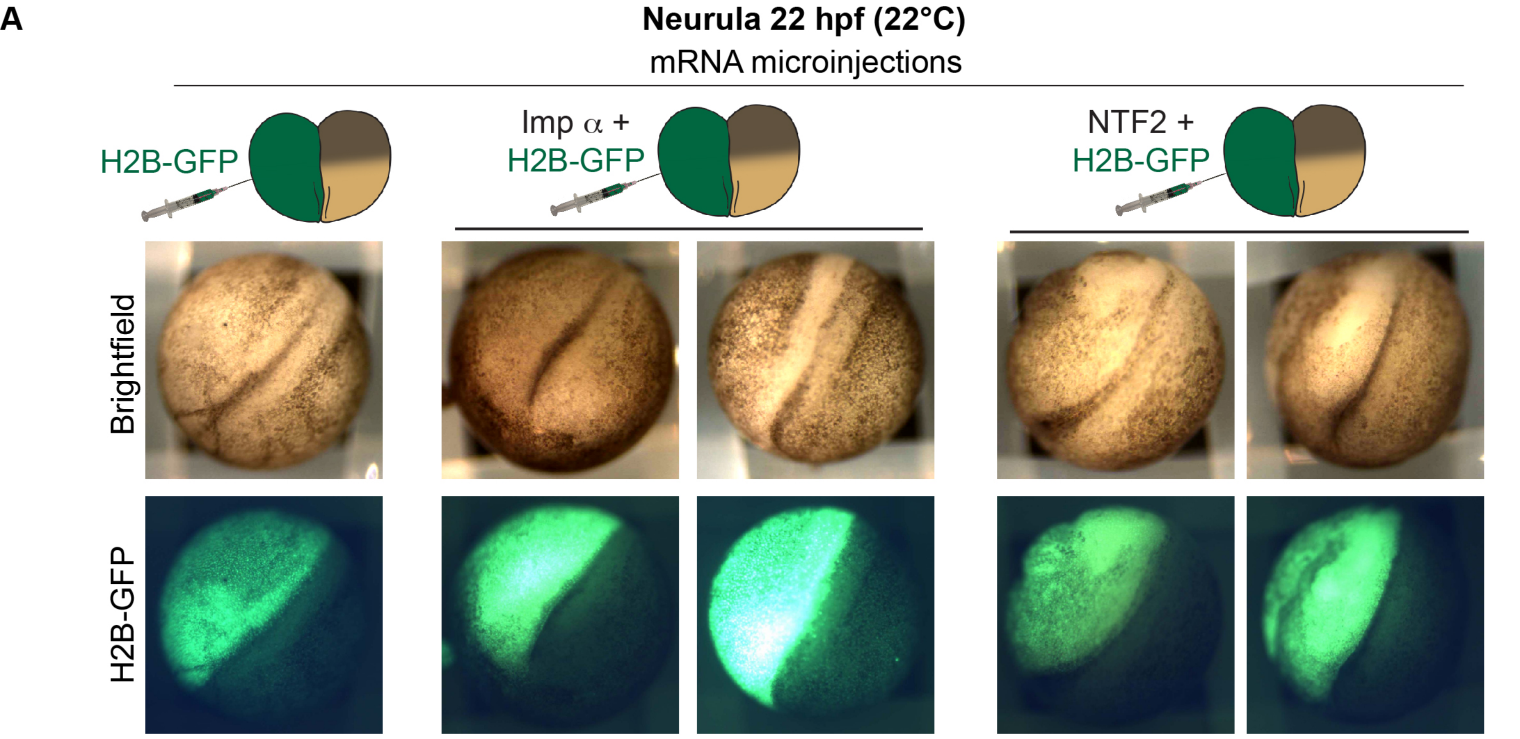
A
Neurula 22 hpf (22°C)
mRNA microinjections
H2B-GFP
Imp α +
H2B-GFP
NTF2 +
H2B-GFP
Brightfield
H2B-GFP

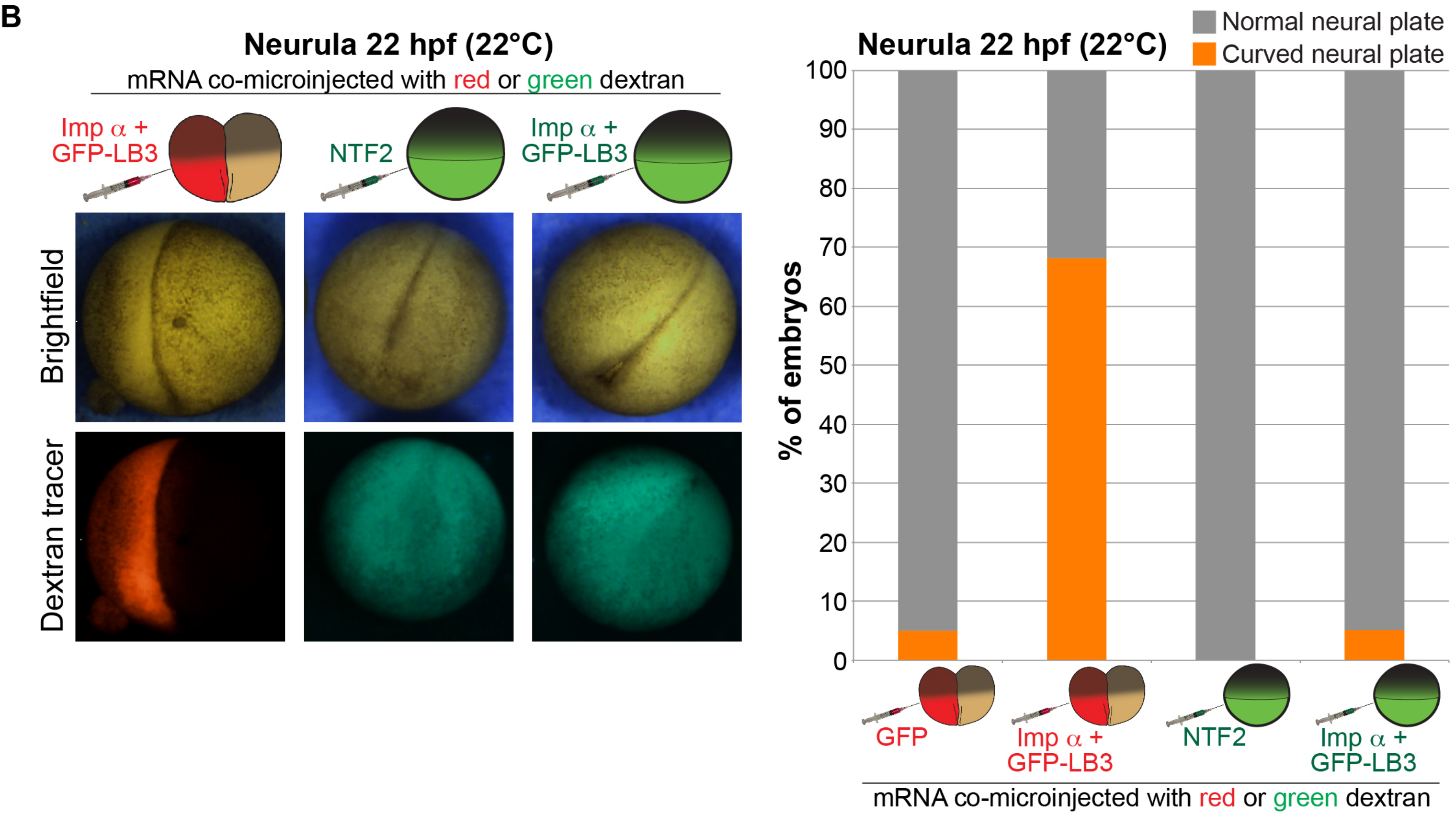
B
Neurula 22 hpf (22°C)
mRNA co-microinjected with red or green dextran
Imp α +
GFP-LB3
NTF2
Imp α +
GFP-LB3
Brightfield
Dextran tracer
Neurula 22 hpf (22°C)
Normal neural plate
Curved neural plate
% of embryos
100
90
80
70
60
50
40
30
20
10
0
GFP
Imp α +
GFP-LB3
NTF2
Imp α +
GFP-LB3
mRNA co-microinjected with red or green dextran

Figure S4

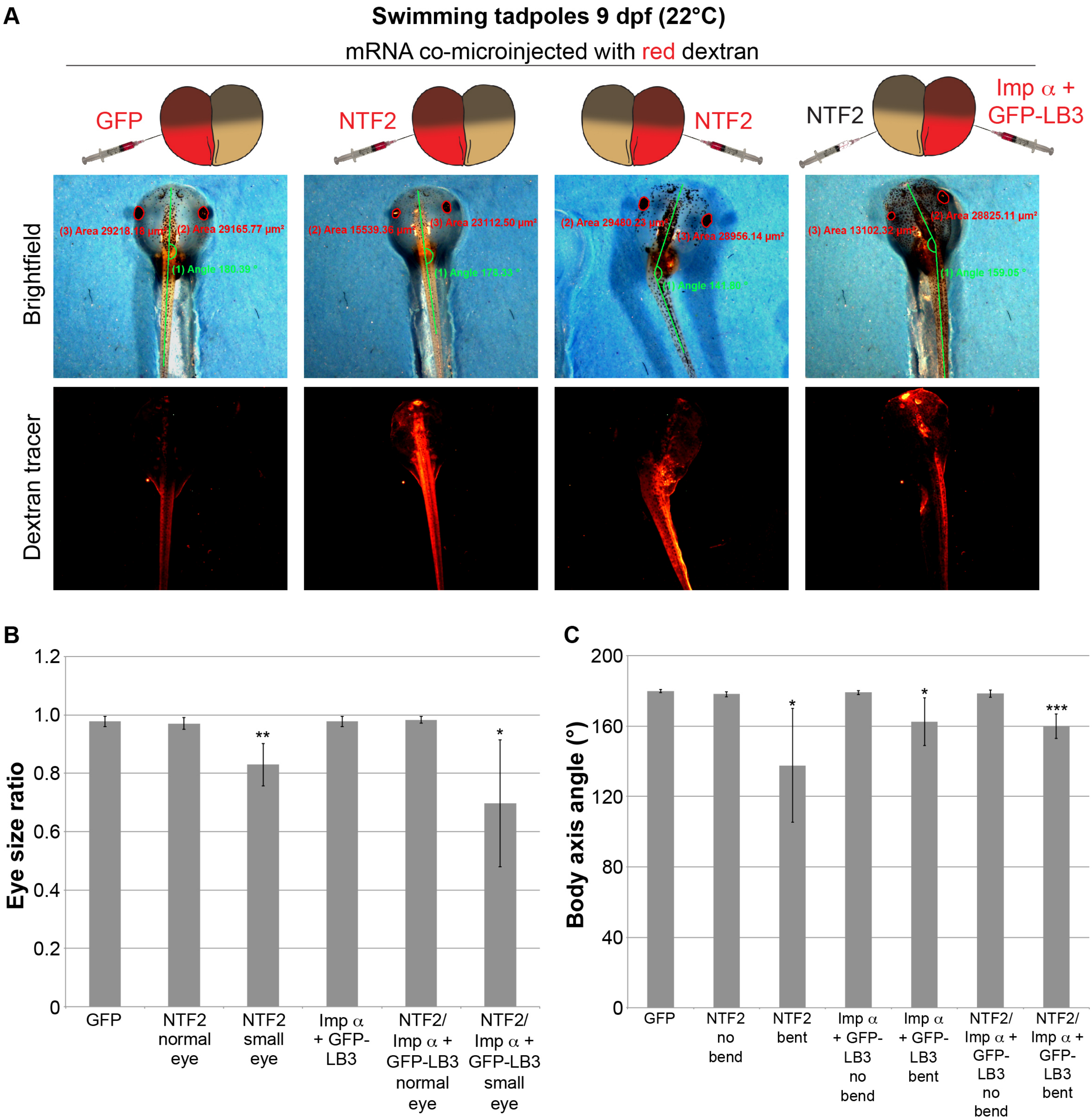
A
Swimming tadpoles 9 dpf (22°C)
mRNA co-microinjected with red dextran
GFP
NTF2
NTF2
NTF2
Imp α + GFP-LB3
Brightfield
Dextran tracer
(3) Area 29218.18 µm²
(2) Area 29165.77 µm²
(1) Angle 180.39 °
(2) Area 15539.36 µm²
(3) Area 23112.50 µm²
(1) Angle 178.33 °
(2) Area 29480.23 µm²
(3) Area 28956.14 µm²
(1) Angle 141.80 °
(3) Area 13102.32 µm²
(2) Area 28825.11 µm²
(1) Angle 159.05 °
B
Eye size ratio
1.2
1.0
0.8
0.6
0.4
0.2
0
**
*
GFP
NTF2 normal eye
NTF2 small eye
Imp α + GFP-LB3
NTF2/ Imp α + GFP-LB3 normal eye
NTF2/ Imp α + GFP-LB3 small eye
C
Body axis angle (°)
200
160
120
80
40
0
*
*
***
GFP
NTF2 no bend
NTF2 bent
Imp α + GFP-LB3 no bend
Imp α + GFP-LB3 bent
NTF2/ Imp α + GFP-LB3 no bend
NTF2/ Imp α + GFP-LB3 bent

Figure S5

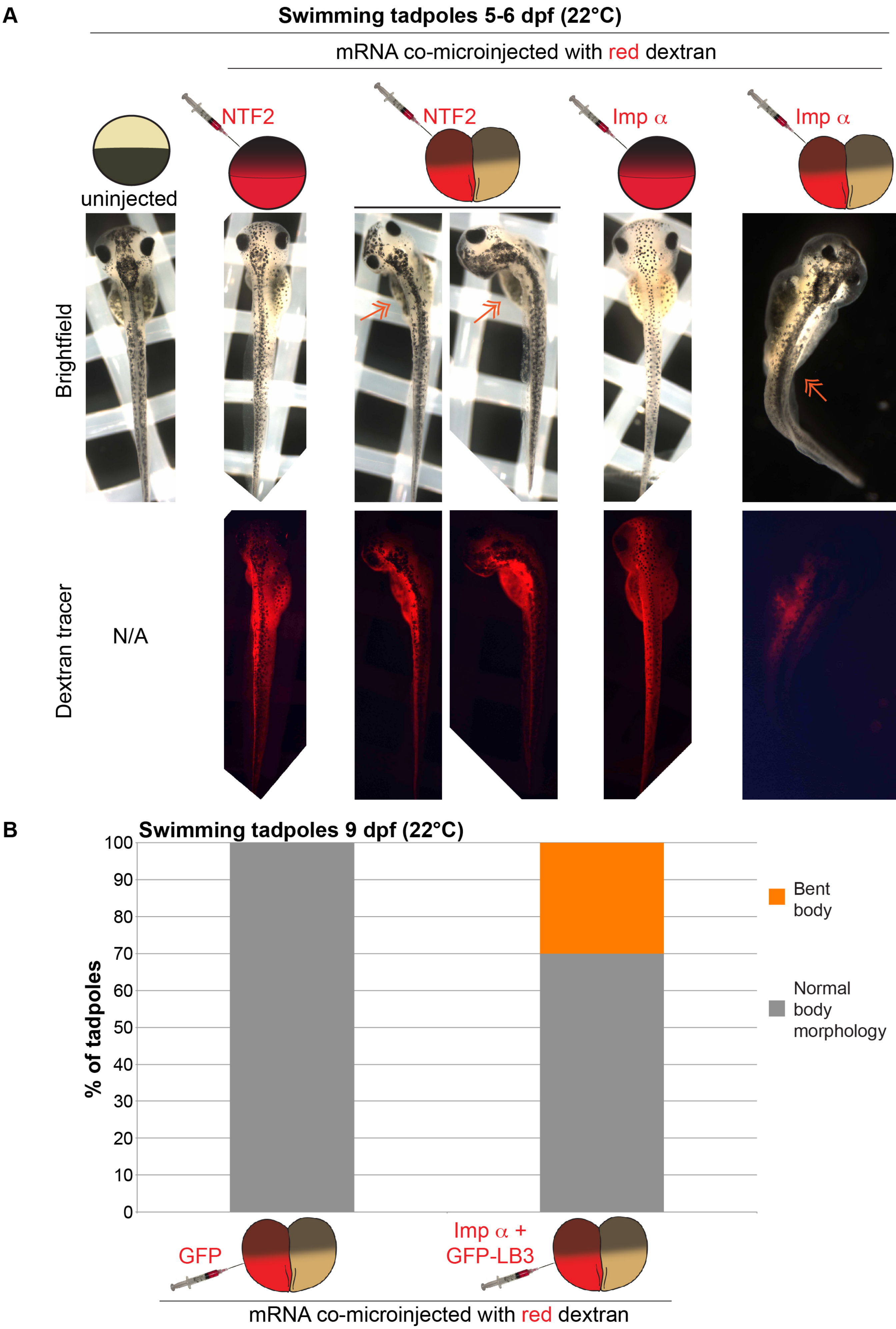
A
Swimming tadpoles 5-6 dpf (22°C)
mRNA co-microinjected with red dextran
uninjected
NTF2
NTF2
Imp α
Imp α
Brightfield
Dextran tracer
N/A
B
Swimming tadpoles 9 dpf (22°C)
% of tadpoles
100
90
80
70
60
50
40
30
20
10
0
Bent body
Normal body morphology
GFP
Imp α + GFP-LB3
mRNA co-microinjected with red dextran

### Figure S6

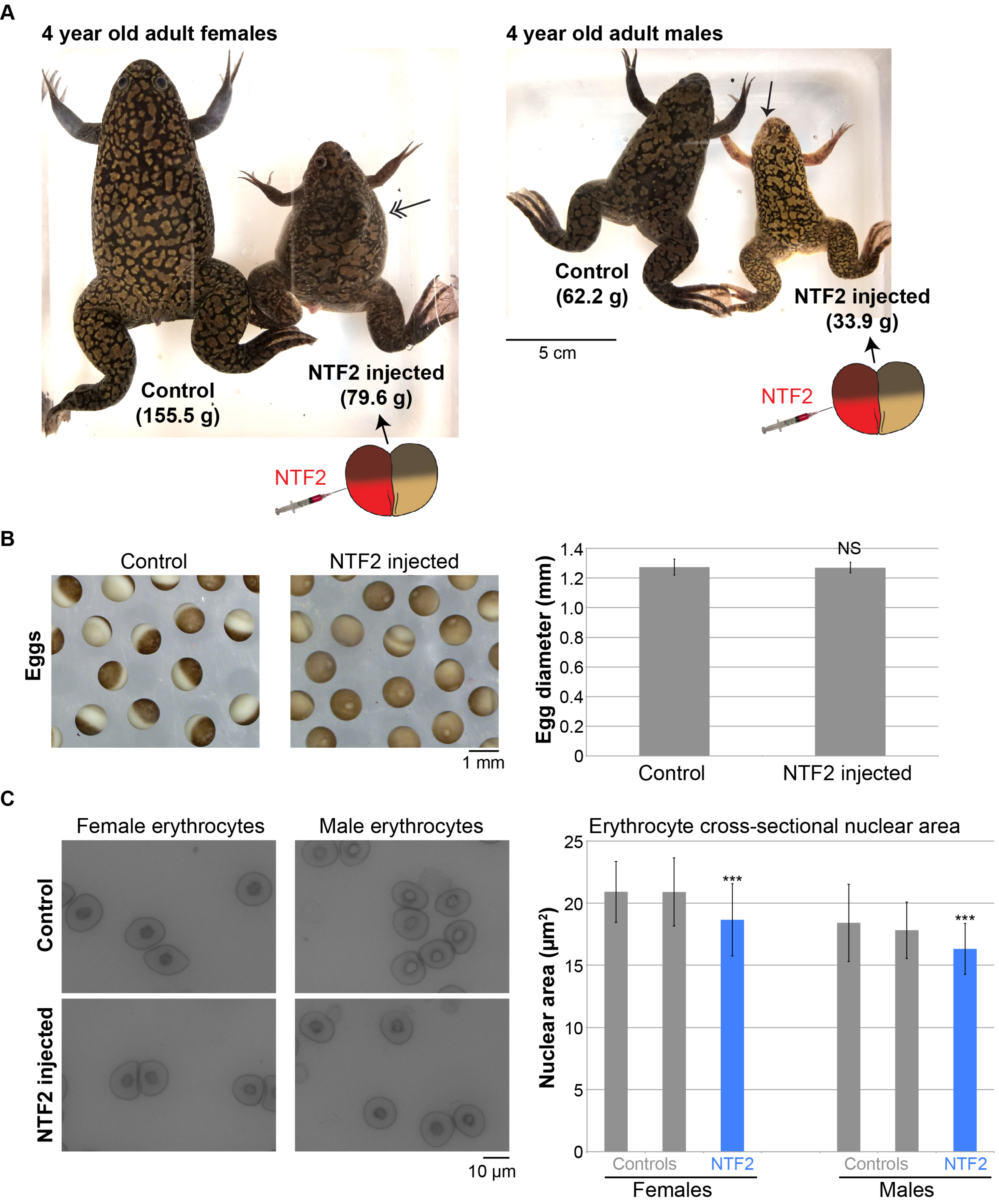
A
4 year old adult females
Control
(155.5 g)
NTF2 injected
(79.6 g)
NTF2
4 year old adult males
Control
(62.2 g)
NTF2 injected
(33.9 g)
5 cm
NTF2
B
Control
NTF2 injected
Eggs
1 mm
Egg diameter (mm)
1.4
1.2
1.0
0.8
0.6
0.4
0.2
0
NS
Control
NTF2 injected
C
Female erythrocytes
Male erythrocytes
Control
NTF2 injected
10 µm
Erythrocyte cross-sectional nuclear area
Nuclear area (µm²)
25
20
15
10
5
0
***
***
Controls
NTF2
Controls
NTF2
Females
Males
